# Involvement of Ca_V_2.2 channels and α_2_δ-1 in hippocampal homeostatic synaptic plasticity

**DOI:** 10.1101/2022.06.27.497782

**Authors:** Kjara S Pilch, Krishma H Ramgoolam, Annette C Dolphin

## Abstract

In the mammalian brain, presynaptic Ca_V_2 channels play a pivotal role for synaptic transmission by mediating fast neurotransmitter exocytosis via influx of Ca^2+^ into the active zone of presynaptic terminals. However, the distribution and modulation of Ca_V_2.2 channels at highly plastic hippocampal synapses remains to be elucidated. Here, we assess Ca_V_2.2 channels during homeostatic synaptic plasticity, a compensatory form of homeostatic control preventing excessive or insufficient neuronal activity during which extensive active zone remodelling has been described. We show that chronic silencing of neuronal activity in mature hippocampal cultures resulted in elevated presynaptic Ca^2+^ transients, mediated by increased levels of Ca_V_2.2 channels at the presynaptic site. This work focussed further on the role of α_2_δ-1 subunits, important regulators of synaptic transmission and Ca_V_2.2 channel abundance at the presynaptic membrane. We find that α_2_δ-1-overexpression reduces the contribution of Ca_V_2.2 channels to total Ca^2+^ flux without altering the amplitude of the Ca^2+^ transients. Levels of endogenous α_2_δ-1 decreased during homeostatic synaptic plasticity, whereas the overexpression of α_2_δ-1 prevented homeostatic synaptic plasticity in hippocampal neurons. Together, this study reveals a key role for Ca_V_2.2 channels and novel roles for α_2_δ-1 during synaptic plastic adaptation.

## Introduction

Synaptic communication relies on the translation of electrical signals to neurotransmitter release^1^. For this, Ca^2+^ enters the presynaptic active zone via voltage-gated Ca^2+^ (Ca_V_) channels, ultimately resulting in the release of synaptic vesicles. This process is tightly regulated by a multitude of proteins but must also remain dynamic to allow rapid adaptation to changes in signalling, such as synaptic potentiation or depression. While excessive or inadequate neuronal firing could impair neuronal activity and brain function, homeostatic synaptic plasticity (HSP) processes maintain firing rates in a physiological range^2^. HSP mechanisms involve changes at both pre- and postsynaptic sites, such as direct regulation of synaptic inputs^3^ and alteration of intrinsic neuronal excitability^4^, for example via presynaptic Ca_V_ channels.

In the mammalian brain, both Ca_V_2.1 (P/Q-type) and Ca_V_2.2 (N-type) channels, each with distinct biophysical properties, provide the main sources of Ca^2+^ influx at the presynapse^5^. The distribution of each subtype varies depending on age, brain region, presynaptic action potential duration and synaptic activity, with some presynaptic terminals exclusively expressing Ca_V_2.1 or Ca_V_2.2^5,6^. Most synaptic transmission, however, is likely mediated by the joint activity of Ca_V_2.1 and Ca_V_2.2 channels, enabling a diversification and fine-tuning of synaptic signalling at central synapses^7^. Due to the relationship between the number of Ca_V_2 channels at the active zone and vesicular release, Ca_V_2 channels play a central role in homeostatic synaptic adaptations.

To study HSP, the sodium channel inhibitor tetrodotoxin (TTX) is widely used to induce chronic silencing of neuronal networks *in vitro*^4^. The prolonged application of TTX has been shown to increase presynaptic Ca^2+^ flux^8^, release probability^9^, and induce restructuring of the active zone, involving multiple presynaptic proteins^10,11^. Among these, presynaptic Ca_V_2.1 channels were identified using live cell Ca^2+^ imaging in rat hippocampal neurons^10,12^. Ca_V_2.2 channels were also found enriched at the presynaptic active zone upon TTX treatment in rat hippocampal neurons^10^. Another study in hippocampal neurons revealed increased presynaptic neurotransmitter release after chronic activity suppression with TTX via Cyclin Dependent Kinase 5/calcineurin modulation of Ca_V_2.2 channels^13,14^.

Ca_V_2 channels rely on auxiliary subunits, particularly α_2_δ, serving as a checkpoint for trafficking and activation of Ca_V_ channels^15,16^. Of the four different α_2_δ isoforms, α_2_δ-1 is of particular interest due to its ubiquitous expression in the brain^17,18^. Moreover, α_2_δ-1 plays a key role for multiple presynaptic^19–21^ and postsynaptic functions^22,23^, potentially independent from its association with Ca_V_ channels^24^.

Here, we show that Ca_V_2.2 channels are involved in HSP in the hippocampus and that the overexpression of α_2_δ-1 affects potentiation of HSP. First, we show that Ca_V_2.2 channels are highly expressed in both young and adult mouse hippocampus and cortex. Second, we demonstrate that TTX treatment increases levels of Ca_V_2.2 channels at presynaptic boutons and their contribution to elevated Ca^2+^ flux in more mature hippocampal neurons. Third, overexpression of α_2_δ-1 results in a downregulation of Ca_V_2.2 channel contribution to basal Ca^2+^ transients. Finally, levels of endogenous α_2_δ-1 decreased during HSP whereas the overexpression of α_2_δ-1 prevented HSP in neurons. Together, these data suggest a key role for Ca_V_2.2 channels and novel roles for α_2_δ-1 during plastic adaptations of synapses.

## Results

### Ca_v_2.2 channels in the immature and mature brain

In the mammalian brain, both Ca_V_2.1 and Ca_V_2.2 channels provide the main sources of Ca^2+^ influx at presynaptic terminals. The abundance of Ca_V_2 channels at the presynapse depends on specific synapse needs and therefore varies between different ages, brain regions, synaptic type and activity^5^.

To determine the relative expression levels of Ca_V_2.2 channels in different brain regions at three different ages, qPCR experiments were performed (Fig 1 A-C). Analysis of mRNA expression in cortex, hippocampus, cerebellum and brain stem revealed similar expression levels of Ca_V_2.2 mRNA in young brains at P0/1 and P7 (apart from lower levels in the brain stem compared to the cortex at P7), whereas in the adult brain, the expression of Ca_V_2.2 channel mRNA was significantly higher in cortex and hippocampus compared to cerebellum and brain stem (Fig 1 A-C, one-way ANOVA, Bonferroni post-hoc). In parallel, to correlate Ca_V_2.2 mRNA levels to endogenous Ca_V_2.2 protein expression, brains from transgenic knock-in mice with an exofacial HA tag on Ca_V_2.2 channels (Ca_V_2.2_HA^KI/KI^;^25^) were used for quantitative immunoblotting of synaptsosomes (Fig 1 D-E). Figure S1 shows immunoblotting of endogenous Ca_V_2.2_HA channels around the somata of hippocampal CA1 cells from Ca_V_2.2_HA^KI/KI^ mice. Immunoblots from brain synaptosomes show that the distribution of Ca_V_2.2 channels follows a trend similar to mRNA levels. At P1 and P7 the distribution of Ca_V_2.2 channels does not vary much between the different regions, whereas in the adult brain, levels of Ca_V_2.2 channels are higher in cortex and hippocampus compared to cerebellum and brain stem (relative to 1 in cortex; levels were 0.89 ± 0.10 in hippocampus, 0.09 ± 0.04 in cerebellum and 0.14 ± 0.06 in brain stem; Fig 1 D-I).

**Figure 1.**
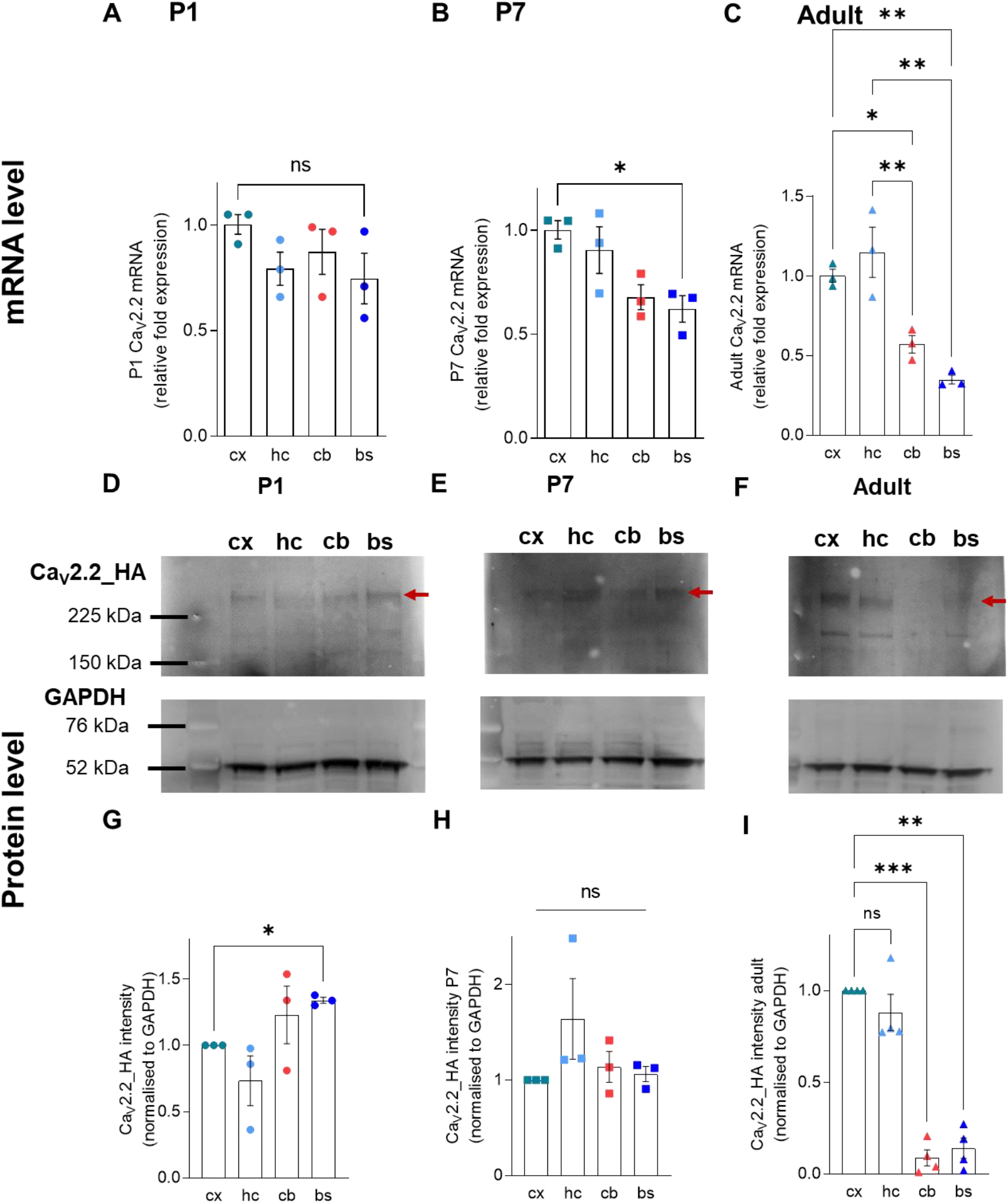
Differential expression of Ca_v_2.2 mRNA and protein levels in different brain regions with most Ca_V_2.2 in the adult cortex and hippocampus. **(A)** At P0/1, mRNA levels of Ca_V_2.2 were smiliar in all four brain regions (cortex (cx) hippocampus (hc), cerebellum (cb) and brain stem (bs)). one-way ANOVA, Bonferroni post-hoc, *F* (3,6) = 1.98 ; *P* = 0.18; n = 3 independent experiments with brains from two pups for each experiment. **(B)** At P7, qPCR experiments reveal higher expression levels of Ca_V_2.2 mRNA in the cortex compared to the brain stem. one-way ANOVA, Bonferroni post-hoc, cx vs bs *F* (3, 6) = 2.3; *P* = 0.02; n = 3 independent experiments with brains from two pups for each experiment. **(C)** In adult brains, Ca_V_2.2 mRNA levels were significantly reduced in cerebellum and brainstem compared to cortex and hippocampus. n = 3 independent experiments. Each n in A-C was assayed in triplicates. All fold changes are made relative to Ca_V_2.2 mRNA levels in the cortex and respective to HPRT mRNA levels and analysed using one-way ANOVA, Bonferroni post-hoc, *F* (3, 6) = 25.40, *** = *P* < 0.001; ** = *P* < 0.01; * = *P* < 0.05 ; ns = not significant. **(D-F)** Immunoblots of synaptosomes from Ca_V_2.2_HA^KI/KI^ mice showing the expression of Ca_V_2.2_HA (red arrow) (top) and GAPDH (bottom) for P0/1 (left), P7 (middle) and adult mice (right) The molecular mass of Ca_V_2.2_HA is 261.0 ± 1.2 kDa, determined from molecular weight markers. **(G)** Western blots showing similar levels of Ca_V_2.2_HA in cortex, hippocampus, cerebellum and brain stem from P0/1 Ca_V_2.2_HA^KI/KI^ mice, with higher levels in brain stem compared to cortex. one-way ANOVA, Bonferroni post-hoc, *F* (1.1, 2.2) = 2.73 ; *P* = 0.23; n = 3 independent experiments with 8 mice pooled for each set. **(H)** Similar levels of Ca_V_2.2_HA in cortex, hippocampus, cerebellum and brain stem from P7 Ca_V_2.2_HA^KI/KI^ mice. one-way ANOVA, Bonferroni post-hoc test, *F* (1, 2.1) = 1.44 ; *P* = 0.35; n = 3 independent experiments with 4 mice pooled for each set. **(I)** In adult mice, Ca_V_2.2_HA levels are higher in the cortex and hippocampus compared to cerebellum and brain stem. Graphs show values from 3 independent experiments. one-way ANOVA, Bonferroni post-hoc test, *F* (1.6, 4.9) = 77.18 ; *P* < 0.001, n = 3 independent experiments, 4 for 12 weeks. All values are normalised to cortex Ca_V_2.2_HA and to respective GAPDH levels; *** = *P* < 0.001; ** = *P* < 0.01; * = *P* < 0.05.

### Induction of homeostatic synaptic plasticity in mature hippocampal neurons

Silencing neuronal activity has been shown to induce compensatory changes at both the pre- and postsynapse^2^. To specifically examine changes in presynaptic Ca^2+^ transients, crucial for neurotransmitter release and therefore presynaptic strength, hippocampal neurons were transfected with genetically encoded Ca^2+^ indicator GCaMP6f coupled to presynaptic synaptophysin (Sy-GCaMP6f^16^; Fig 2). Neurons were stimulated with 1 AP (Fig 2 A), and then 10 AP (Fig 2 B) to identify responding boutons. Co-expression of pH indicator mOr2, coupled to synaptic vesicle associated membrane protein (VAMP), allowed for targeted analysis of Ca^2+^ transients at neurotransmitter-releasing synaptic boutons (Fig 2 C)^26^. For analysis of Ca^2+^ transients, only the responses from releasing (orange line, Fig 2 F) boutons were included. Fig 2 D and E show changes in fluorescence in *ΔF/F*_*0*_ of releasing (open circles) and of non-releasing (closed circles) boutons after stimulation with 1 (Fig 2 D) and 10 (Fig 2 E) APs.

**Figure 2.**
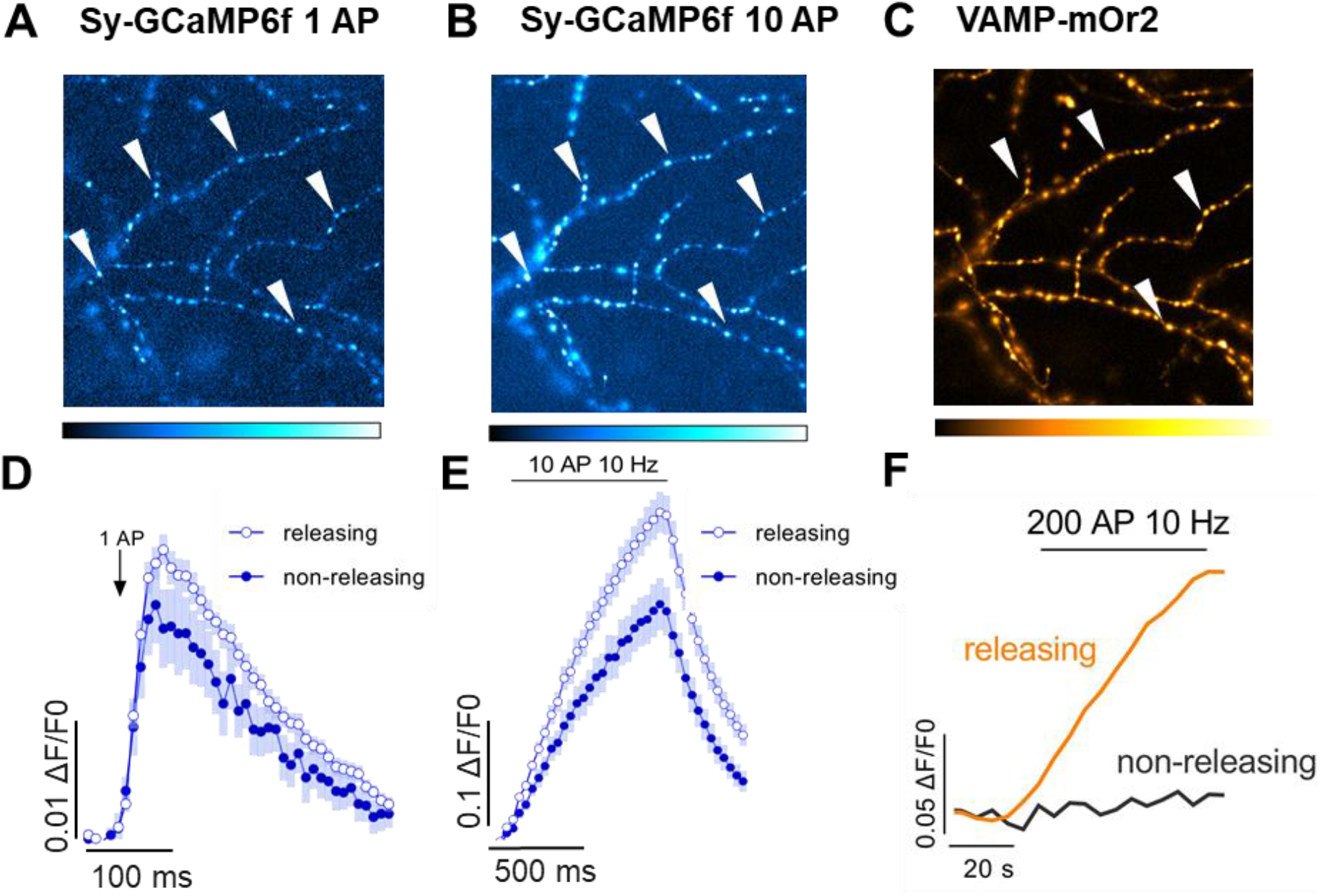
Monitoring presynaptic Ca^2+^ transients in hippocampal neurons with Sy-GCaMP6f and VAMP-mOr2. **(A, B)** Images showing the expression of Sy-GCaMP6f in putative boutons during stimulation with 1 AP (arrowheads). **(B)** Changes in Sy-GCaMP6f fluorescence after stimulation with 10 APs were used to identify responding boutons (arrowheads). **(C)** VAMP-mOr2 fluorescence after stimulation with 200 APs at 10 Hz was used to identify functional vesicle-releasing synapses based on an increase in fluorescence following the increase in pH to which it is exposed during vesicle fusion. **(D)** Up to 75 ROIs were selected per field of view to show changes in fluorescence over baseline *(ΔF/F0)* of averages of 5-8 repeats of 1 AP stimulation. Responses from releasing (blue open circles) and non-releasing (blue filled circles) boutons were distinguished based on VAMP-mOr2 responses. **(E)** Responses from releasing (blue open circles) and non-releasing (blue filled circles) boutons after stimulation with 10 APs. **(F)** Increases in VAMP-mOr2 fluorescence (orange line) after stimulation with 200 APs was used to identify releasing boutons (orange line) and non-releasing boutons (black line). Based on this, responses to 1 and 10 AP were categorized into releasing and non-releasing boutons.

To induce homeostatic changes at the synapse, neurons were incubated with TTX for 48 h and Ca^2+^ transients were measured at two ages: between day in vitro (DIV) 14 and 17 and between DIV 18 and 22 (Fig 3). In younger cultures, DIV 14-17, no significant changes in Ca^2+^ transient amplitudes were detected between control and TTX-treated neurons (0.032 ± 0.003 in control cells and 0.04 ± 0.005 after TTX treatment, Fig 3 A and B). However, in more mature cultures at DIV 18-22, measured Ca^2+^ transient amplitudes were much larger following TTX treatment (0.024 ± 0.002 in control neurons and 0.038 ± 0.002 in TTX-treated neurons, Fig 3 C and D).

**Figure 3.**
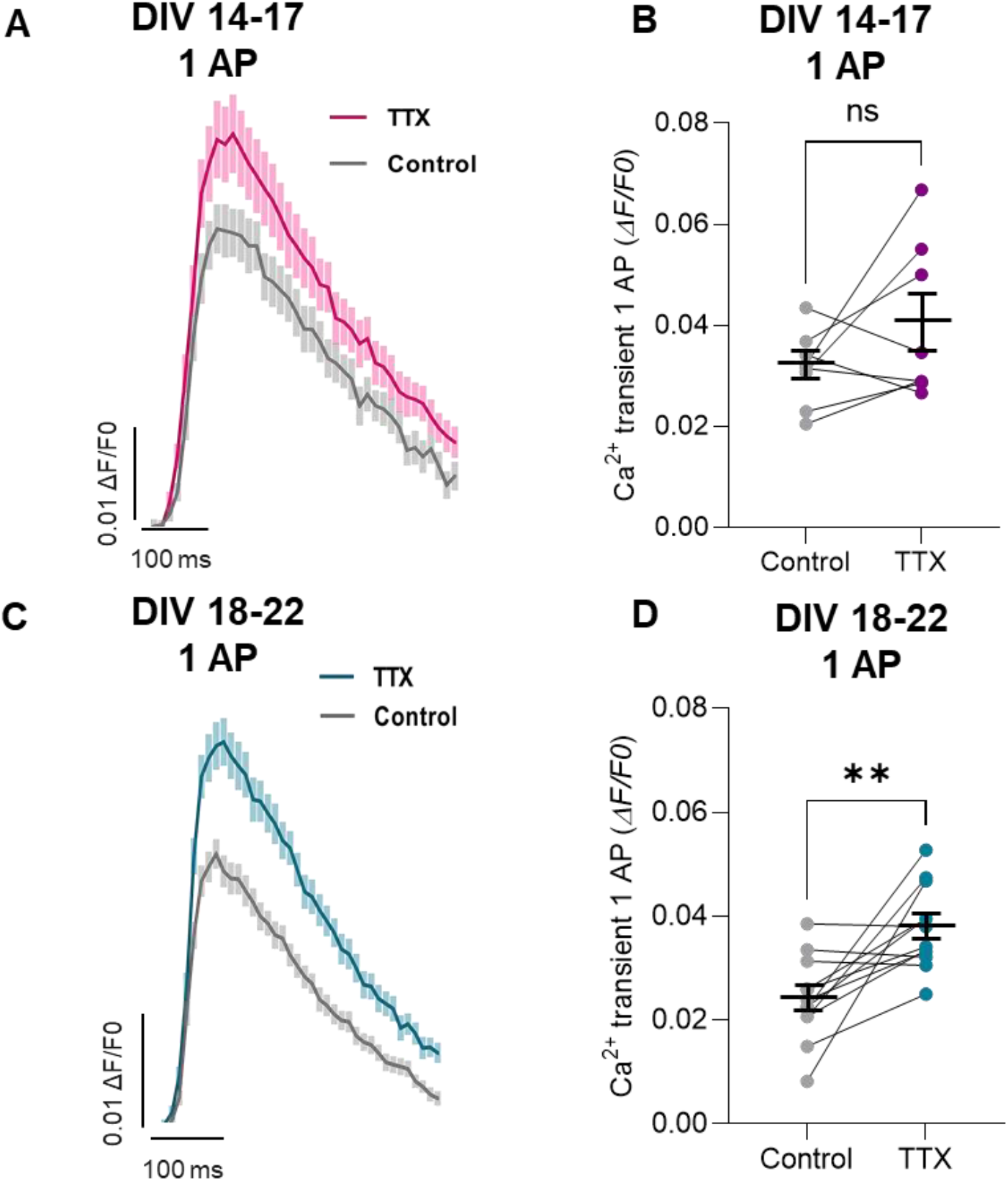
TTX treatment increases presynaptic Ca^2+^ transient amplitudes in mature mouse hippocampal boutons. **(A)** Sy-GCaMP6f fluorescence changes in functionally-releasing presynaptic boutons after stimulation with 1 AP of control (grey) and TTX (magenta) conditions at DIV 14-17. Averaged fluorescence traces of 5-8 repeats of electrical stimulation with 1 AP. For each field of view, 10-75 ROIs were chosen for analysis of change in fluorescence over baseline fluorescence *(ΔF/F0)*. **(B)** At DIV 14-17, no significant differences were observed between presynaptic Ca^2+^ transient amplitudes in control (grey) and TTX-treated (magenta) boutons, n = 8 biological replicates, two-tailed paired t-test *P* = 0.16; t = 1.56; df = 7; n corresponds to independent experiments and data are also shown as mean ± SEM (black). **(C)** Sy-GCaMP6f fluorescence changes at DIV 18-22 in TTX (blue) treated presynaptic terminals and control (grey) untreated terminals. Averaged traces of 5-8 repeats of electrical stimulation with 1 AP. For each field of view, 10-75 ROIs were chosen for analysis of change in fluorescence over baseline fluorescence *(ΔF/F0)*. **(D)** At DIV 18-22, TTX treatment (blue) induced an increase in presynaptic Ca^2+^ transient amplitudes compared to control boutons (grey), n = 12 biological replicates, two-tailed paired t-test, *P* = 0.003; t = 3.72; df = 22; n corresponds to independent experiments and data are also shown as mean ± SEM (black).

### Increased levels of Ca_v_2.2 channels contribute to larger Ca^2+^ transients during HSP

At presynaptic terminals, a combination of different Ca_V_2 channels mediate Ca^2+^ influx. To examine the contribution of Ca_V_2.2 channels during HSP, cells were stimulated before and after the application of the Ca_V_2.2 channel-specific inhibitor ω-conotoxin GVIA (ConTx). At DIV 14-17, Ca_V_2.2 channel contribution to Ca^2+^ transients during 1 AP was similar in control and TTX-treated neurons. In control and TTX-treated neurons, 31.4 ± 7.4 % and 46.8 ± 5.9 % of total Ca^2+^ influx was mediated by Ca_V_2.2 channels, respectively (Fig 4 A). Conversely, during HSP, in more mature neurons, the contribution of Ca_V_2.2 channels to overall Ca^2+^ influx increased. At DIV 18-22 the contribution of Ca_V_2.2 to Ca^2+^ flux rose from 28.6 ± 8.4 % in control neurons to 54.94 ± 4.7 % in potentiated hippocampal neurons treated with TTX (Fig 4 B). Moreover, immunoblotting of WCL of hippocampal neurons at DIV 18-22 showed a 37.6 ± 0.02 % increase in level of Ca_V_2.2 channels after the induction of HSP with TTX (Fig 4 C and D). To determine whether this increase was associated with preysnaptic boutons, we used hippocampal neurons cultured from transgenic Ca_V_2.2_HA^KI/KI^ mice to visualise endogenous Ca_V_2.2 channels *in vitro* (Fig 4 E). Quantification of Ca_V_2.2_HA intensity associated with puncta positive for the presynaptic marker vGluT1 revealed increased levels of Ca_V_2.2_HA channels in neurons treated with TTX (control normalized mean intensity 100 ± 8.4 % and TTX mean intensity 127.2 ± 10.5 %; Fig 4 F and G). Figure S2 shows the specificity of the Ca_V_2.2_HA staining as no HA-signal is detectable in neurons from wild-type mice.

**Figure 4.**
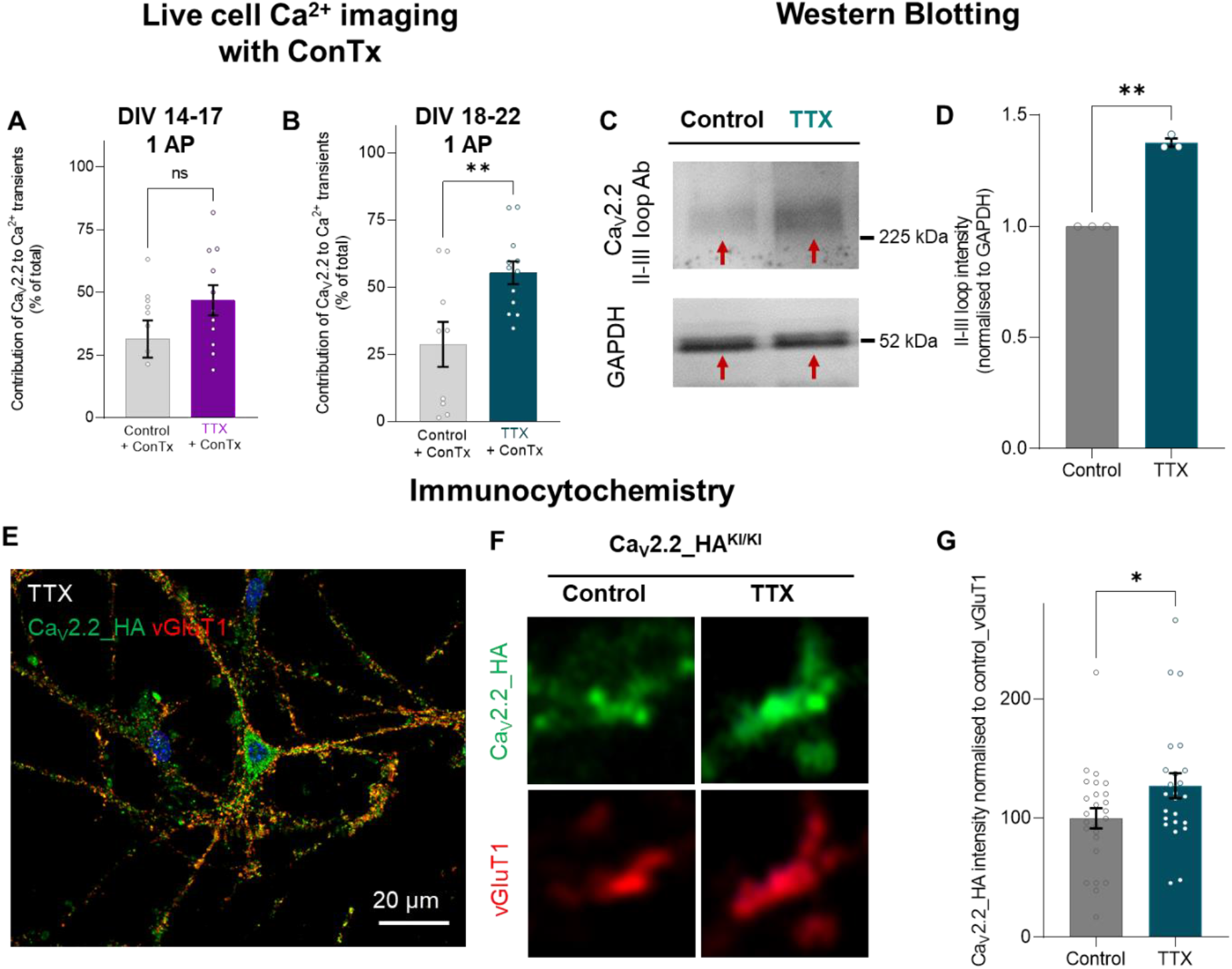
Ca_V_2.2 channels mediate increased Ca^2+^ transients during HSP at DIV 18-22. **(A)** At DIV 14-17, contribution of Ca_V_2.2 channels to Ca^2+^ transient amplitudes after electrical stimulation with 1 AP is similar to untreated neurons (grey bar) and after TTX incubation (purple bar and data). Two-tailed unpaired t-test, *P* = 0.12; t = 1.63; df = 19; n for control = 10 and n for TTX = 11. n corresponds to fields of view from 3 independent experiments and is shown as mean ± SEM. **(B)** At DIV 18-22, Ca_V_2.2 channels contribute more to Ca^2+^ flux after the induction of HSP with TTX. Two-tailed unpaired t-test *P* = 0.01; t = 3.05; df = 19; n for control = 9 and n for TTX = 11. n numbers correspond to fields of view from 3 independent experiments and is shown as mean ± SEM. **(C)** Representative immunoblots using Abs against the II-III loop of Ca_V_2.2 channels in WCL of control neurons and TTX-treated neurons (red arrowhead, top gel) at DIV 18-22. Values were normalised to the intensity of GAPDH for control and TTX neurons, respectively (red arrow heads, bottom gel). **(D)** Quantification of the intensity of Ca_V_2.2 II-III loop band of 3 independent experiments reveals a stronger signal after the induction of HSP with TTX (blue column) compared to control, untreated neurons (grey column). Two-tailed paired t-test *P* = 0.002; t = 20.2; df = 2; n = 3 independent experiments, averaged duplicates for each experiment. **(E)** Airyscan image of TTX-treated hippocampal neuron from Ca_V_2.2_HA^KI/KI^ mouse at DIV 21 stained with anti-HA Abs (green) and anti-vGluT1 Abs (red) to identify presynaptic terminals. Magnification x 63, scale bar 20 μm. **(F)** 3 × 3 μm subset images from control boutons (left panels) and TTX-treated (right panels) with Ca_V_2.2_HA (green) and vGluT1 (red). **(G)** Bar chart showing increased Ca_V_2.2_HA intensity in TTX-treated neurons compared to control neurons as mean ± SEM. n control neurons = 25 and n TTX-treated = 24 neurons, from 3 independent experiments. For each neuron, up to 75 ROIs were selected. Data were normalised to the averaged control value. Two-tailed unpaired t-test, *P* = 0.048; t = 2.23; df = 49.

### α_2_δ-1 overexpression does not change Ca^2+^ transients resulting from 1 AP stimulation and prevents HSP

The Ca_V_ α_2_δ subunits are emerging as an important regulators of presynaptic organisation and function^15,24,27^. Hence, we investigated the effect of α_2_δ-1-overexpression on Ca^2+^ transient amplitudes and the Ca_V_2.2 channel contribution to Ca^2+^ flux (Fig 5).

**Figure 5.**
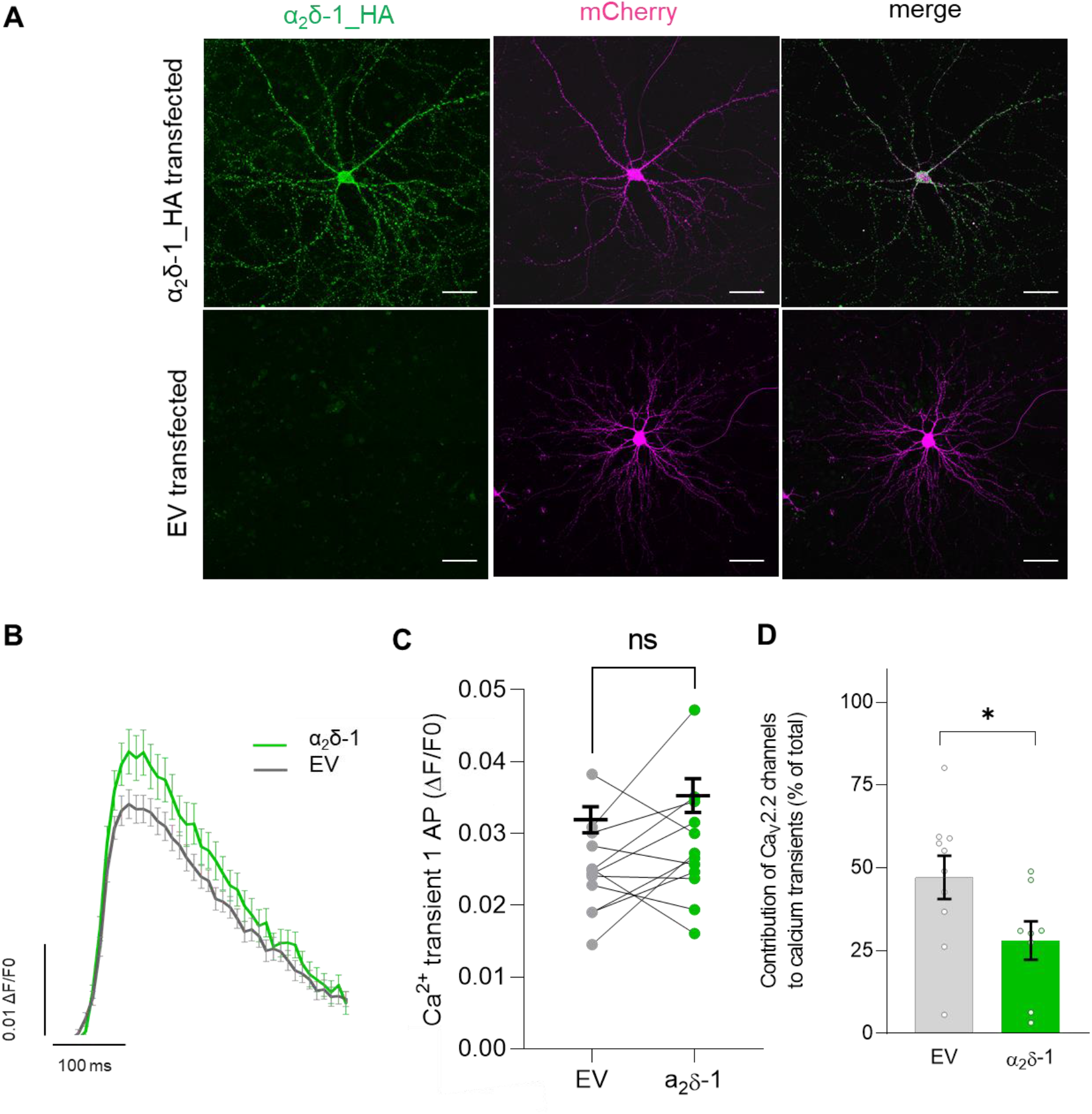
Overexpression of α_2_δ-1 does not change Ca^2+^ transients resulting from 1 AP, but decreases the contribution of Ca_V_2.2 channels. **(A)** Confocal images of immunostaining for α_2_δ-1_HA in α_2_δ-1 overexpressing neurons (top row) and control, EV-transfected neurons (bottom row). α_2_δ-1_HA is shown in green and transfection marker mCherry in magenta. Maximum intensity projection of z-stacks and tile scan, optical section 0.279 μm, confocal mode x 40, scale bar = 50 μm. **(B)** Similar fluorescence changes in releasing Sy-GCaMP6f expressing boutons over time after stimulation with 1 AP of control EV (grey line) and α_2_δ-1-overexpressing (green line) neurons. Averaged fluorescence traces of 5-8 repeats of stimulation with 1 AP. For each field of view, 10-75 ROIs were chosen for analysis of change in fluorescence over baseline fluorescence (*ΔF/F0*). **(C)** Ca^2+^ transient amplitudes after stimulation with 1 AP were similar in EV (grey) and α_2_δ-1-overexpressing (green) neurons, n = 12 biological replicates, paired t-test *P* = 0.18, t = 1.53, df = 11. n corresponds to independent experiments and data are also shown as mean ± SEM (black). **(D)** Contribution of Ca_V_2.2 channels to Ca^2+^ transients after stimulation with 1 AP was 42.1 ± 6.4 % (grey) in EV neurons and 28.1 ± 5.8 % (green) in neurons overexpressing α_2_δ-1, n for EV control = 10 fields of view and n for α_2_δ-1 = 8 fields of view. n corresponds to fields of view from 5 independent experiments and is shown as mean ± SEM. Two-tailed unpaired t-test, *P* = 0.005; t = 2.13; df = 16.

The overexpression of α_2_δ-1 is shown in Fig 5 A using anti-HA Abs against tagged α_2_δ-1-HA; no background staining is visible in the EV control neurons. Comparison of Ca^2+^ transients after 1 AP stimulation showed no changes in Ca^2+^ influx (Fig 5 B and C) with similar fluorescence profiles in the EV control (grey, 0.025 ± 0.002) and the α_2_δ-1-overexpressing boutons (green, 0.028 ± 0.002). Since we found no changes in Ca^2+^ transient amplitudes in α_2_δ-1-overexpressing neurons, we then applied ConTx to assess if the contribtion of Ca_V_2.2 channels to Ca^2+^ transients was altered. ConTx application revealed that in α_2_δ-1-overexpressing neurons, the contribution of Ca_V_2.2 channels to Ca^2+^ transients after 1 AP stimulation is decreased to 28.1 ± 5.8 % compared to 42.1 ± 6.4 % in EV-transfected neurons (Fig 5 D). Figure S3 shows the control immunostaining for anti-α_2_δ-1 Abs.

The α_2_δ-1 subunit is important not only for the trafficking and function of Ca_V_2.2 channels, but it is also emerging as an important transsynaptic regulator with several described functions independent from its association with Ca_V_2 channels^24^. To elucidate if the induction of HSP changes the surface distribution of endogenous α_2_δ-1, biotinylation assays were performed using control and TTX-treated neurons at DIV 18-22 (Fig 6 A, B). The hippocampal neurons potentiated with TTX revealed a decrease of surface (biotinylated fraction) and total endogenous α_2_δ-1 (WCL fraction) (decrease to 0.88 ± 0.01 and to 0.86 ± 0.04, respectively; Fig. 6 A and B). To then further assess the role of α_2_δ-1 for HSP, we overexpressed α_2_δ-1 in neurons and recorded Ca^2+^ transients after incubation with TTX (Fig 6 C). In neurons overexpressing α_2_δ-1, no increases in Sy-GCaMP6f Ca^2+^ transient amplitudes were induced by treatment with TTX, whereas Ca^2+^ transients in EV untreated control neurons were larger after TTX treatment, similar to values shown in Fig 3 (0.023 ± 0.001 in EV untreated neurons and 0.043 ± 0.005 in TTX-treated EV neurons; 0.027 ± 0.001 in α_2_δ-1-overexpressing neurons and 0.029 ± 0.003 in TTX-treated α_2_δ-1 overexpressing neurons, Fig. 6 C). These findings could have important implications for the role of α_2_δ-1 in regulating both Ca_V_2.2 channel plasticity and other mechanisms involved in HSP.

**Figure 6.**
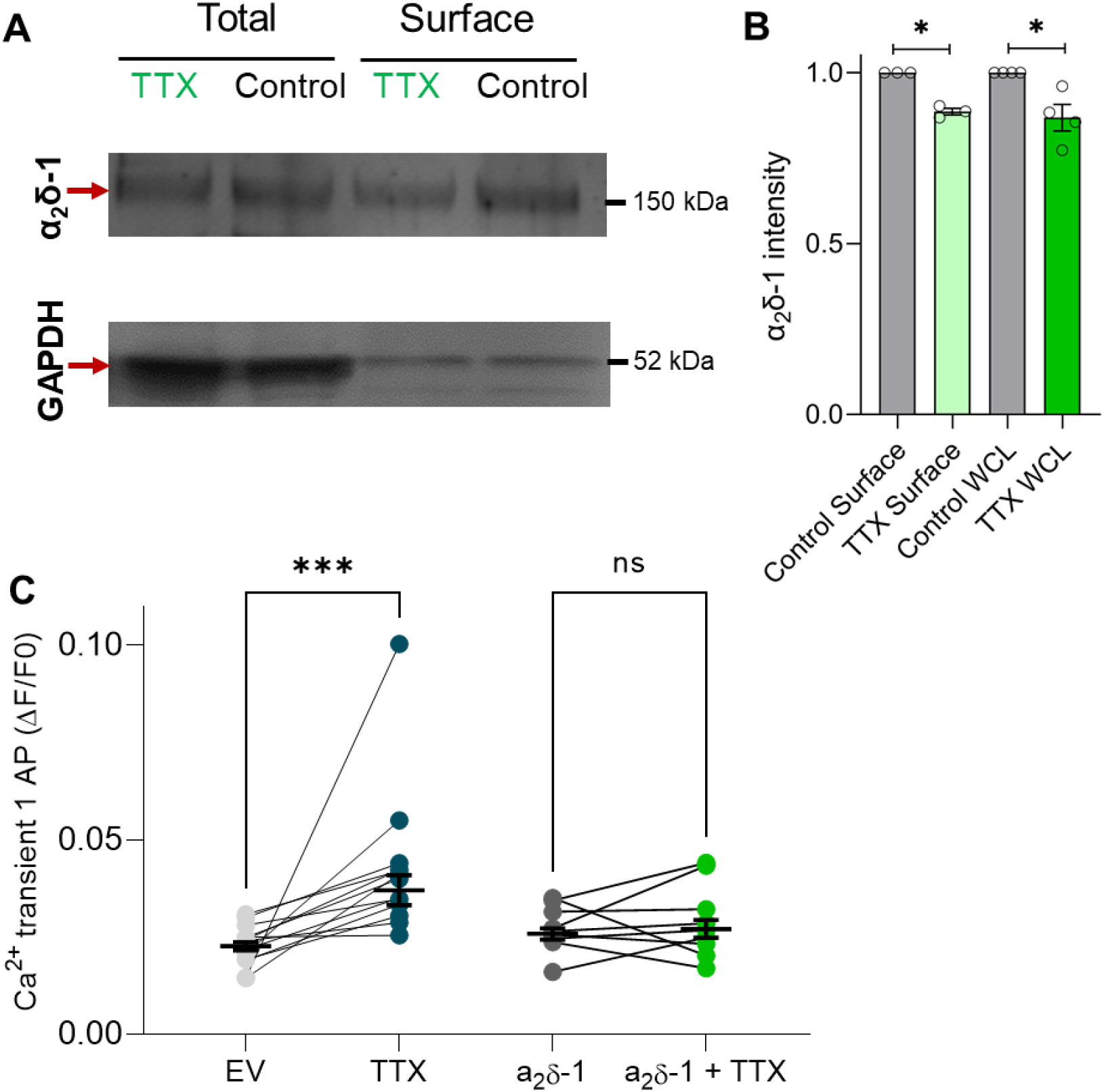
During HSP, surface and total levels of endogenous α_2_δ-1 decrease while overexpression of α_2_δ-1 prevents HSP. **(A)** Biotinylation experiments of control and TTX-treated hippocampal neurons showing the total endogenous amount of α_2_δ-1 (WCL) and surface fractions, respectively (top gel, red arrows). Bands for GAPDH are shown in the bottom gel (red arrows). **(B)** Quantification of the intensity of endogenous α_2_δ-1 protein bands of 3 independent experiments reveal decreased levels of total and surface α_2_δ-1 after the induction of HSP with TTX (green columns) compared to control, untreated neurons (grey columns). Paired t-test *P* = 0.02, n = 3 independent experiments, averaged duplicates for each experiment. **(C)** Ca^2+^ transient amplitudes after stimulation with 1 AP are larger in TTX-treated EV than in control EV conditions (grey data points) but similar in α_2_δ-1 overexpressing neurons with (green) and without TTX (dark grey data points). One-way ANOVA, Tukey’s multiple comparisons test, *F* (3,40) = 1.3; EV vs EV + TTX *P* < 0.001; α_2_δ-1 vs α_2_δ-1 + TTX *P* = 0.1; n corresponds to independent experiments and data are shown as mean ± sem. n for EV and EV + TTX = 13 and n for α_2_δ-1 and α_2_δ-1 + TTX = 9.

## Discussion

Presynaptic Ca_V_2 channels play a pivotal role in synaptic transmission by mediating fast neurotransmitter exocytosis via influx of Ca^2+^ into the active zone. Here, we combine gene expression studies, immunoblotting, immunocyochemistry and live cell Ca^2+^ imaging to show i) the role of Ca_V_2.2 channels for synaptic transmission in the hippocampus, ii) upregulation of Ca_V_2.2 channels mediates increased Ca^2+^ flux during HSP, iii) HSP downregulates endogenous α_2_δ-1 subunits at hippocampal synapses and iv) overexpression of α_2_δ-1 decreases the contribution of Ca_V_2.2 to presynaptic Ca^2+^ flux and abolishes the effect of TTX on elevated Ca^2+^ transients.

### Ca_v_2.2 channels in the adult hippocampus

We show high levels of Ca_V_2.2 channel protein and mRNA expression in mouse cortex and hippocampus throughout development, persisting into adulthood. While younger mice showed relatively similar expression of Ca_V_2.2 mRNA levels across brain regions, adult cortex and hippocampus had higher levels compared to cerebellum and brain stem. This finding was supported by protein immunoblot studies using synaptosomal fractionation of brains from Ca_V_2.2_HA^KI/KI^ mice.

One of the key questions regarding Ca_V_2.2 channels is how they are distributed and regulated in different brain regions and at different synapses. Initial work on this examined the Calyx of Held, a large auditory relay synapse in the brain stem. Before hearing onset in young mice, synaptic transmission is mediated by loosely coupled Ca_V_2.1 and Ca_V_2.2 channels, whereas in the mature synapse Ca^2+^ flux is mediated by tightly coupled Ca_V_2.1 channels^28–30^. This nanodomain coupling of Ca_V_2.1 channels allows rapid and temporally precise glutamate release, required for auditory processing. Although the developmental shift from Ca_V_2.1 to Ca_V_2.2 channels has also been described in the neocortex^31^, hippocampus^32^ and cerebellum^33^, our findings indicate that this developmental down-regulation of Ca_V_2.2 channels may not relate to the majority of cortical and hippocampal synapses. Indeed, there is evidence that Ca_V_2.2 channels at least partially mediate presynaptic Ca^2+^ flux in the adult cortex and hippocampus^15,26,34–37^. Highly plastic synapses, such as in the hippocampus, may use a combination of Ca_V_2.1, Ca_V_2.2 and Ca_V_2.3 channels, each providing the synapse with distinct coupling properties, activity-dependent facilitation and modulation of Ca_V_2 channels. This may enable dynamic changes in synaptic output depending on synaptic activity^5,38^. The hippocampal mossy fiber pathway, for example, was shown to rely on microdomain coupling of Ca_V_2.2 channels for presynaptic plasticity^39^. Notably, the shape of APs in different brain regions and neuron types (e.g. narrow APs in interneurons and in the Calyx of Held versus broader APs at the hippocampal presynapse^6^) might be another factor determining which Ca_V_2 channel subtype is predominantly activated. The mechanisms underlying the differential distribution of Ca_V_2.1 and Ca_V_2.2 channels remain unclear.

### Increased levels of Ca_v_2.2 channels contribute to increased Ca^2+^ flux during HSP

Our observation that about 30 % of Ca^2+^ influx occurs via Ca_V_2.2 channels in mature mouse hippocampal neurons is similar to previous findings^35,36^, which is suggestive of an important role for Ca_V_2.2 channels in synaptic transmission in mature neurons. Chronic silencing of neuronal activity with TTX in hippocampal cultures resulted in larger presynaptic Ca^2+^ transients, as already described in previous studies^10,12^. In the present study, elevated presynaptic Ca^2+^ transients were exclusively observed in more mature cultures (18-22 DIV) following incubation with TTX. This is in line with previous findings of HSP adaptations being mostly postsynaptic in younger neurons, whereas presynaptic adaptations emerge as neurons mature^40,41^.

Presynaptic HSP involves dynamic restructuring of the active zone matrix^11^ and a recruitment of proteins of the presynaptic machinery^10^, including Ca_V_2.1 channels^10,12^. Our findings indicate that an increased contribution of Ca_V_2.2 channels to presynaptic Ca^2+^ flux (from about 30 % to 50 %, Fig 4 B, D and G.) represents another component of presynaptic HSP restructuring in cultured hippocampal neurons. This is further confirmed by both western blotting and immunocytochemistry of Ca_V_2.2_HA^KI/KI^ neurons, revealing increased levels of Ca_V_2.2 channels following HSP induction of approximately 25-30 %. The increase in Ca_V_2.2 channel expression during HSP has not been previously described, though a recent study detected an enrichment of Cav2.2 channels at the active zone after TTX treatment using STORM super resolution imaging^10^. The precise details of presynaptic restructuring during HSP, as well as the relevance of the change in relative composition of active zone Ca_V_2 channels remain to be fully deciphered.

### α_2_δ-1 overexpression downregulates Ca_v_2.2 channel involvement in Ca^2+^ response to 1 AP stimulation and prevents HSP

α_2_δ-1 subunits are important parameters in synaptic transmission, by regulating Ca_V_2 channel abundance at the presynaptic membrane^15^. In addition, α_2_δ-1 potentially interacts with other proteins to modulate synaptic activity, independent from Ca_V_2 channels^24,42^. Here, we show that α_2_δ-1-overexpression reduces the contribution of Ca_V_2.2 channels to total Ca^2+^ flux without altering the amplitude of the Ca^2+^ transients. In contrast, previous work showed that α_2_δ-1 overexpression reduced Ca^2+^ flux relative to synaptic vesicular release^15^. One possible explanation for our findings is that in more mature cultures, the downregulation of Ca_V_2.2 channel plasticity as a result of α_2_δ-1 overexpression was accompanied by a compensatory upregulation of other Ca_V_2 channels, ensuring stable Ca^2+^ transients.

Generally, the amount of α_2_δ proteins is likely to vary in different synapses, but levels are usually thought to be higher than those of Ca_V_2 channels^43^. It was shown that the complex between α_2_δ and Ca_V_2 channels is easily disrupted^43,44^, hence the more α_2_δ is present, the more Ca_V_2 channels will be in complex with α_2_δ. α_2_δ proteins are glycosyl-phosphatidylinositol-anchored^45^ and are present in lipids rafts, which are small microdomains within the plasma membrane, high in cholesterol and sphingolipids (for review see^46^). Notably, α_2_δ-1 also mediates the partitioning of Ca_V_2.1 and Ca_V_2.2 channels into these specialised lipid-rich membrane domains^47,48^, resulting in reduced Ca^2+^ currents^47,49^. Increasing the abundance of α_2_δ-1 in presynaptic terminals by overexpression may lead to increased levels of Ca_V_2.2 channels in lipid rafts, potentially clamping their mobility, and their contribution to the plasticity of Ca^2+^ flux. The fact that Ca^2+^ transients in response to 1 AP did not change in α_2_δ-1-overexpressing terminals may be due to compensatory Ca_V_2.1 or Ca_V_2.3 channel upregulation, ensuring stable Ca^2+^ transients. Further work is needed to evaluate the stability of the interaction between α_2_δ-1 and Ca_V_2 channels at the different compartments of the active zone.

Furthermore, α_2_δ-1 overexpression prevents elevated presynaptic Ca^2+^ transients observed after TTX treatment. A recent study investigated the effect of α_2_δ-1 overexpression on the developmental of neuronal networks *in vitro* and discovered that neurons overexpressing α_2_δ-1 exhibit spontaneous neuronal activity and increased presynaptic glutamate release^50^. This finding might indicate that neuronal network activity in α_2_δ-1 overexpressing neurons was already increased, therefore the application of TTX did not have a potentiating effect. This increased neuronal activity might also explain the downregulation of Ca_V_2.2 channels. We therefore also sought to determine changes in endogenous surface α_2_δ-1 during HSP by cell surface biotinylation and immunoblotting. This revealed lower levels of α_2_δ-1 both in WCL, and specifically on the surface of neurons after the induction of HSP with TTX, indicating that a downregulation of α_2_δ-1 may contribute to processes involved in HSP.

Chronic silencing of neuronal activity thus caused an increase in Ca^2+^ transients due to greater Ca_V_2.2 channel contribution and increased Ca_V_2.2 protein levels, while levels of α_2_δ-1 decreased. This decrease of α_2_δ-1 may therefore be required to allow increased mobility of Ca_V_2.2 channels, necessary for presynaptic potentiation. Overexpression of α_2_δ-1 potentially prevents the elevation of presynaptic Ca^2+^ transients after TTX treatment by “clamping” Ca_V_2.2 channels in microdomains within the plasma membrane.

Together, these findings show an involvement of Ca_V_2.2 channels in HSP and prompt further examination into the role of α_2_δ proteins, as major regulators of homeostatic processes at synapses.

## Materials and Methods

### Animals

Mice were housed in groups of no more than five on a 12-h:12-h light:dark cycle; food and water were available *ad libitum*. All experimental procedures were covered by UK Home Office licenses, had local ethical approval by University College London (UCL) Bloomsbury Animal Welfare and Ethical Review Body.

### qPCR

For gene expression studies, brains from postnatal (P) 0/1, P7 and 12 weeks old mice were separated into cortex, hippocampus, cerebellum and brain stem and disrupted using a rotor-stator homogeniser (Disperser T10, IKA, Staufen, Germany). Total RNA was extracted using RNeasy lipid tissue mini kit according to manufacturer’s instructions. RNA concentrations were photometrically measured to reversely transcribe 5 μg RNA from each sample into complementary DNA using High-Capacity RNA-to-cDNA kit. For the 50-cycle qPCR (1 h at 37 °C, 5 min at 95 °C), triplicates from each sample (3 different mice for each timepoint) were loaded into a 96-well plate with TaqMan Universal PCR Master Mix (Applied Biosystems) and the following TaqMan probes used (gene name: assay ID): Hypoxanthine-phosphoribosyltransferase 1 (*Hprt1*): Mm00446968m1; *Cacna1b*: Mm01333678m1. Optimal threshold values were defined automatically as 0.1 by 7500 Real-Time PCR software (Applied Biosystems) and used to determine the cycle threshold number (C_T_). Results are expressed as fold change in Ca_V_2.2 mRNA expression, given as mean ± sem. Data were normalised to expression levels of internal control gene *hprt* and analysed using the 2^-ΔΔC^_T_ method^51^. To ensure sufficient amounts of RNA at timepoint P0/1 and P7, brains from 2 mice were pooled. For each age, 3 independent RNA extractions were performed and run on the same plate.

### Subcellular fractionation

Brains from P0/1, P7 and 12 weeks old Ca_V_2.2_HA^KI/KI^ mice^25^ were dissected in buffer containing 0.32 M sucrose, 3 mM HEPES, 0.25 mM dithiothreitol, pH 7.4 and cOmplete protease inhibitor cocktail (Roche) and separated into cortex, hippocampus, brain stem and cerebellum. Synaptosomes were prepared as previously described^52^. Crude synaptosomes were solubilised for 30 min on ice in 50 mM Tris, 150 mM NaCl, 1 % Igepal, 0.5 % Na deoxycholate, 0.1 % sodium dodecyl sulfate (SDS), cOmplete protease inhibitor cocktail (Roche), pH 8^53^. After centrifugation, protein concentrations were determined using the Bradford protein assay method (BioRad). Samples were adjusted to the same concentration and denaturated with 100 mM dithiothreitol reducing agent and Laemmli sample buffer at 55 °C for 10 min. 20 μg of protein were loaded onto a 3-8 % NuPAGE gel and proteins resolved by SDS-polyacrylamide gel electrophoresis (SDS-PAGE) for 1 h 05 min at 150 V, 50 mA in running buffer. Following the separation of proteins, they were transferred from the gel to a polyvinylidene difluoride (PVDF) membrane using a semi-dry transfer blot (BioRad) for 10 min at 25 mV, 1 mA. The membrane was then blocked by incubation with 3 % bovine serum albumin (BSA), 10 mM Tris pH 7.4, 0.5 % Igepal for one h. Following an overnight incubation at 4 °C with rat anti-HA (1:500, monoclonal, Roche, Cat # 11867423001) or mouse anti-GAPDH (1:25.000, polyclonal, Ambion, Cat # AM4300) antibodies (Abs), membranes were incubated for one h at room temperature (RT) with horseradish peroxidase (HRP)-coupled secondary Abs at 1:2000 for one h (all secondary Abs from Biorad, raised in goatanti-rat HRP Cat # 5204-2504 and anti-mouse HRP Cat # 1721011). Protein bands were revealed using ECL reagent (ECL 2, Thermo Scientific) with a Typhoon 9419 phosphorimager (GE Healthcare) and analysed using ImageJ software.

### Whole-cell lysate (WCL) immunoblotting

WCL for immunoblotting were prepared by transferring control and TTX-treated neurons at DIV 18-22 on ice, washing them twice with PBS containing 1 mM CaCl_2_ and MgCl_2_ and collecting cells by scraping them in CaCl_2_/MgCl_2_ PBS containing cOmplete protease inhibitor cocktail (Roche). Lysates were cleared by centrifuging at 1000 x g for 10 min at 4 °C and pellets were resuspended in PBS with 1 % Igepal, 0.1 % SDS, 0.5 % sodium deoxycholate and protease inhibitor. After brief sonication and rotation for one h at 4 °C, cells centrifuged at 16000 x g for 30 min at 4 °C. The protein concentration was determined using Bradford protein assay (Biorad) and were run on a 3-8 % NuPAGE gel as described above. The PVDF membrane was cut according to weight amrkers to be able to use the mouse Abs twice, then blocked and incubated overnight at 4 °C with rabbit anti-Ca_v_2.2 II-III loop Abs at 1:500, mouse anti-α_2_δ-1 Abs (polyclonal, Roche, Cat # C5105) at 1:1000 or with mouse anti-GAPDH (1:25;000) Abs. After washing with TBS 0.5 % Igepal, membranes were incubated for 1 h at RT with HRP-coupled secondary Abs at 1:2000 for one hour. Anti-rabbit HRP sAbs were Cat # 1706515. Protein bands were revealed as described above.

### Biotinylation immunoblotting

Biotinylation experiments were adapted from our previous paper^16^. Control and TTX-treated hippocampal neurons at DIV 18-22 were washed twice with Hanks Balanced Salt Solution (HBSS) containing 1 mM CaCl_2_ and MgCl_2_ (modified HBSS). They were then incubated with Premium Grade EZ-link Sulfo-NHS-LC-Biotin (Thermo Scientific; 1 mg/ml) in modified HBSS for 30 min at RT. After quenching with 200 mM glycine, cells were transferred to ice, washed twice with modified HBSS and collected by scraping. Following centrifugation at 1000 x g for 10 min at 4 °C, the pellet was frozen at -80 °C until sufficient numbers of samples were collected for pooling (up to 4 independent preps). Afterwards, cells were lysed in lysis buffer containing 1 % Igepal, 0.1% SDS, 0.5 % sodium deoxycholate and protease inhibitor in HBSS, sonicated and rotated for one h at 4 °C. Subsequently, protein concentrations were determined using the Bradford method as described above. Streptavidin beads were then added to a fraction of the cells while keeping some of the sample as a whole cell lysate to run on the same western blot. Streptavidin-treated samples were left on the roller at 4 °C overnight and proteins revealed as decribed above.

### Primary neuronal cultures and transfection

Primary neuronal cultures were prepared from hippocampi of P0/1 wild-type or Ca_V_2.2_HA^KI/KI^ mice^25^. After neck dislocation, hippocampi were dissected in ice-cold dissection medium (HBSS, HEPES 1 M, 1 % w/v glucose) and dissociated in enzyme solution containing Papain solution (HBSS, 2 mg/ml L-Cysteine, 2 mg/ml bovine serum albumin (BSA) 50 mg/ml Glucose), Papain (70 U/ml) and DNase I (1200 U/ml) in HBSS. After digestion at 37 °C for 40 min, the enzyme solution was aspirated and prewarmed inactivation medium (minimum essential media (MEM), 5 % v/v foetal bovine serum (FBS), 0.38 % w/v Glucose, 0.25 % w/v BSA) added. Hippocampi were then triturated with a P1000 micropipette with polypropylene plastic tips and cells centrifuged at RT at 1000 rpm for 10 min in serum medium (MEM, 5 % v/v FBS, 1.38 % w/v glucose). After cell pellet resuspension, cells were counted and plated at a concentration of 6800 cells/mm^2^ on coverslips precoated with poly-D-Lysine (50 μg/ml). Cells were covered with serum-free neuronal plating medium comprising Neurobasal medium, supplemented with B27™ and GlutaMAX™ and kept at 37 °C in 5 % CO_2_ with a medium change every 3 days.

At DIV 7 cells were transfected using Lipofectamine 2000 Transfection reagent according to manufacturer’s instructions. Cells were transfected with synaptophysin-GCaMP6f (Sy-GCaMP6f, (Kadurin et al., 2016) and VAMP-mOrange 2 (VAMP-mOr2) at a ratio of 3:1 ^26^. VAMP-mOrange2 was generated by replacing mCherry from pCAGGs-VAMP-mCherry by mOrange2 (gifts from Tim Ryan), Sy-GCaMP6f was made by replacing GCaMP3 in pCMV-SyGCaMP3 (a gift from Tim Ryan) by GCaMP6f^54^. For experiments with α_2_δ-1 overexpression, cells were transfected with Sy-GCaMPf6, VAMP-mOr2 and either α_2_δ-1 (pCAGGS, rat, genbank accession number M86621) or empty vector (EV) at a ratio of 2:1:1. For immunocytochemistry experiments, cells were transfected with mCherry and α_2_δ-1-HA and EV at 1:1:2 for α_2_δ-1 over expressing cells or with mCherry and EV at 1:3 for control conditions. 2 h prior to transfection, half of the cell media was replaced with fresh media and fresh media was added to the previously removed media to obtain conditioned media. Transfection mixes for one coverslip contained 4 μg DNA in 50 μl OptiMEM and 2 μl Lipofectamine in 50 μl OptiMEM, added dropwise to the cells. After 2 h in the incubator, media was replaced with conditioned media.

### Immunohistochemistry

Immunohistochemical experiments were performed using Ca_V_2.2_HA^KI/KI^ and wild-type mice of 12 weeks age. Mice were anaesthetized with an intraperitoneal injection of pentobarbitone (Euthatal, Merial Animal Health, Harlow, UK; 600 mg / kg), transcardially perfused with saline containing heparin (0.1 M) followed by perfusion with 4 % w/v paraformaldehyde (PFA) in 0.1 M phosphate buffer (pH 7.4) at a flow rate of 2.5 ml/min for 5 min. Following perfusion, brains were postfixed in 4 % PFA for 2 h and immersed in cryoprotective 20 % sucrose overnight. Subsequently, brains were mounted in optimal cutting temperature compound and sliced into 20 μm thick coronal sections using a cryostat and then stored at -80 °C. Slices were blocked and permeabilised for one h at RT in 10 % v/v goat serum (GS) and 0.2 % v/v Triton-X 100 in PBS and then further blocked by applying unconjugated goat F(ab) anti-mouse IgG (1:100) for 1 h at RT to prevent unspecific binding. Subsequently, primary Abs (rat anti-HA (as above) and anti-vGAT, rabbit polyclonal, 1:500, synaptic systems, #131003) were applied diluted in 5 % v/v GS, 0.2 % v/v Triton-X 100 and 0.005 % v/v NaN_3_ overnight at 4 °C. After 2 days, 4 % PFA was reapplied for 30 min to ensure stabilisation of the protein-antibody complex. Sections were then incubated with the goat anti-rat and anti-rabbit secondary Abs conjugated with Alexa Fluor 488 and Alexa Fluor 594, respectively, for 2 days at 4 °C (Invitrogen, both 1:500). After washing, sections were mounted in Vectashield Antifade Mounting Medium (Vector Laboratories). Images were taken using a LSM 780 confocal microscope (Zeiss) with a x20 objective (89 μm optical section, pixel dwell 2.05 μs, zoom 1.8) or x 63 in super resolution mode. After acquisition, super resolution images underwent Airyscan processing and tile stitching using Zen software (Zeiss).

### Immunocytochemistry

For the staining of endogenous Ca_V_2.2_HA channels from Ca_V_2.2_HA^KI/KI^ mice, hippocampal neurons at DIV 18-22 were fixed with 1 % w/v PFA/ 4 % w/v sucrose in PBS for 5 min followed by washes in PBS. Next, neurons were blocked and permeabilised for at least one h at RT in 20 % v/v horse serum, 0.1 % v/v Triton X100. Subsequently, cultures were incubated overnight at 4 °C with primary Abs diluted in in 10 % v/v horse serum, 0.1 % v/v Triton X-100 (rat anti-HA as above and anti-vGluT1 (guinea-pig, polyclonal, 1:1000, synaptic systems). Next, cells were post-fixed with 1 % w/v PFA/ 4 % w/v sucrose in PBS for 5 min at RT and incubated with respective donkey Alexa Fluor secondary Abs applied for 1 h diluted at 1:500. Following washes in PBS, cells were mounted in Vectashield and examined using Confocal or super resolution Airyscan mode imaging on a LSM 780 confocal microscopes (Zeiss) with x 63 objective (1768 × 1768 pixels) as z-stacks (0.173 μm optical section). To quantify the signal intensity of Ca_V_2.2_HA, up to 75 regions of interest of 2 μm in diameter were manually chosen based on vGluT1 and Ca_V_2.2_HA colocalization on a single plane and the intensity measured in ZEN (Zeiss, version 5). Values from TTX boutons were normalised to control values. Analysis was performed blind and randomized.

For the α_2_δ1-HA staining, hippocampal cells were fixed with 4 % w/v PFA/ 4 % w/v sucrose in PBS for 5 min followed by washes in PBS. Next, cells were blocked and permeabilised for at least 1 h at RT in 20 % v/v goat serum, 0.3 % v/v Triton X-100. Subsequently, cultures were incubated overnight at 4 °C with rat anti-HA Abs at 1:200 (Roche) and guinea-pig anti-red fluorescent protein (RFP) Abs at 1:500 (synaptic systems) diluted in 10 % v/v goat serum, 0.3 % v/v Triton X-100 (Ab solution). Next, cells were washed with PBS and goat anti-rat Alexa Fluor 488 and goat anti-guinea-pig Alexa Fluor 594 secondary Abs were applied for one h diluted at 1:500 in Ab solution. After washing with PBS, cells were mounted using Vectashield. Images were acquired using confocal mode of LSM 780 confocal microscopes (Zeiss) with a 40x oil-immersion objective in 8-bit mode. Images were taken as tile (2×2 each consisting of 1024 × 1024 pixels) and z-stack scans (0.28 μm optical section) with a pixel dwell of 2.05 μs.

### Live cell Ca^2+^ imaging

Ca^2+^ imaging experiments were performed as described in a previous paper^26^. Plated cells on 22 mm^2^ coverslips were transferred to a laminar-flow perfusion and stimulation chamber (Imaging chamber with field stimulation series 20, multichannel systems) and mounted on an epifluorescence microscope (Axiovert 200 M, Zeiss) under continuous perfusion at 23 °C at 0.5 ml/min^−1^ with Ca^2+^ perfusion buffer containing (in mM) 119 NaCl, 2.5 KCl, 2 CaCl_2_, 2 MgCl_2_, 25 HEPES (buffered to pH 7.4) and 30 glucose. In order to suppress postsynaptic activity, 10 μM 6-cyano-7-nitroquinoxaline-2,3-dione (CNQX, Sigma) and 50 μM D, L-2-amino-5-phosphonovaleric acid (AP5, Sigma) were included. For some experiments, irreversible Ca_V_2.2 channel inhibitor ω-conotoxin GVIA (ConTx; 1 μM, Alomone) was applied for 2 min before stimulation under continuous perfusion. To ascertain whether any observed reduction in fluorescence was due to the incubation period or due to bleaching during re-stimulation, control experiments were performed with normal imaging medium applied for 2 min instead of ConTx and cells re-stimulated which resulted in a reduction of 8.8 ± 9.0 % during 1 AP stimulation. All values shown have been adjusted for this reduction.

Homeostatic presynaptic plasticity was induced by incubating cells with 1.5 mM tetrodotoxin (TTX) for 48 h prior to imaging^8^. Before imaging, cells were incubated in Ca^2+^ perfusion buffer for 20 min to wash out TTX. Images were acquired with an Andor iXon+ (model DU-897U-CS0-BV) back-illuminated EMCCD camera using OptoMorph software (Cairn Research, UK) with LEDs as light sources (Cairn Research, UK). Fluorescence excitation and collection was done through a 40 × 1.3 NA Fluar Zeiss objective using 450/50 nm excitation and 510/50 nm emission and 480 nm dichroic filters (for Sy-GCaMP6f) and a 572/35 nm excitation and low-pass 590 nm emission and 580 nm dichroic filters (for VAMP-mOr2). APs were evoked by passing 1 ms current pulses via platinum electrodes. Transfected boutons were selected for imaging by stimulating neurons with trains of 6 APs at 33 Hz using Digitimer D4030 and DS2 isolated voltage stimulator (Digitimer Ltd.). To measure calcium responses, neurons were stimulated with a single AP (repeated at least 5 times with 30 s time intervals to improve signal-to noise ratio) and then with 10 APs at 10 Hz. Synaptic boutons were identified by VAMP-mOr2 expression and defined as functional based on responsiveness to 200 APs stimulation at 10 Hz. From each coverslip, a maximum of 3 fields of views were recorded. When ConTx was applied, only one field of view was imaged per coverslip. To get a baseline value of fluorescence, 20 frames were recorded before stimulation (*F*_*0*_). Images were acquired at 100 Hz and 7 ms exposure time and up to 75 putative synaptic boutons within the image field were selected for analysis using a 2 μm region of interest (ROI) and analysed in ImageJ (http://www.rsb.info.nih.gov/ij) using a custom-written plugin (http://www.rsb.info.nih.gov/ij/plugins/time-series.html). Data was background adjusted and changes calculated as change of fluorescence intensity over baseline fluorescence before stimulation (*ΔF/F*_*0*_). For presentational purposes, images were adjusted for brightness and contrast.

### Statistics

Data are given as mean ± sem with the number of independent experiments (n) and statistical test used specified for each figure. Results were considered significant with a *P* value < 0.05. Data were analysed and graphs generated with GraphPad Prism 9.

## Author contributions

Conceptualisation K.S.P and A.C.D. Experiments and analysis K.S.P and K.H.R. Writing K.S.P. Funding acquisition A.C.D. All authors commented on the paper.

## Acknowledgements

We thank Dr Laurent Ferron for his help with initial live cell imaging experiments and Dr Ivan Kadurin for his help with initial western blotting experiments. We thank Wendy S Pratt and Kanchan Chaggar for skillful technical assistance and Stuart Martin for genotyping. We thank Dr Joshua Elliott for help with proof-reading the manuscript. This study was supported by an Investigator award from the Wellcome Trust (grant no. 206279\Z\17\Z to A.C.D.)

