## Supplementary data for "Involvement of Ca_V_2.2 channels and α_2_δ-1 in hippocampal homeostatic synaptic plasticity"

### Involvement of Ca$_V$2.2 channels and $\alpha_2\delta$-1 in hippocampal homeostatic synaptic plasticity

**Figure S1 (relates to Fig 1 D-I). Visualisation of Ca$_V$2.2_HA channels in pyramidal neurons of the CA1 region in the adult hippocampus of Ca$_V$2.2_HA$^{KI/KI}$ mice.**

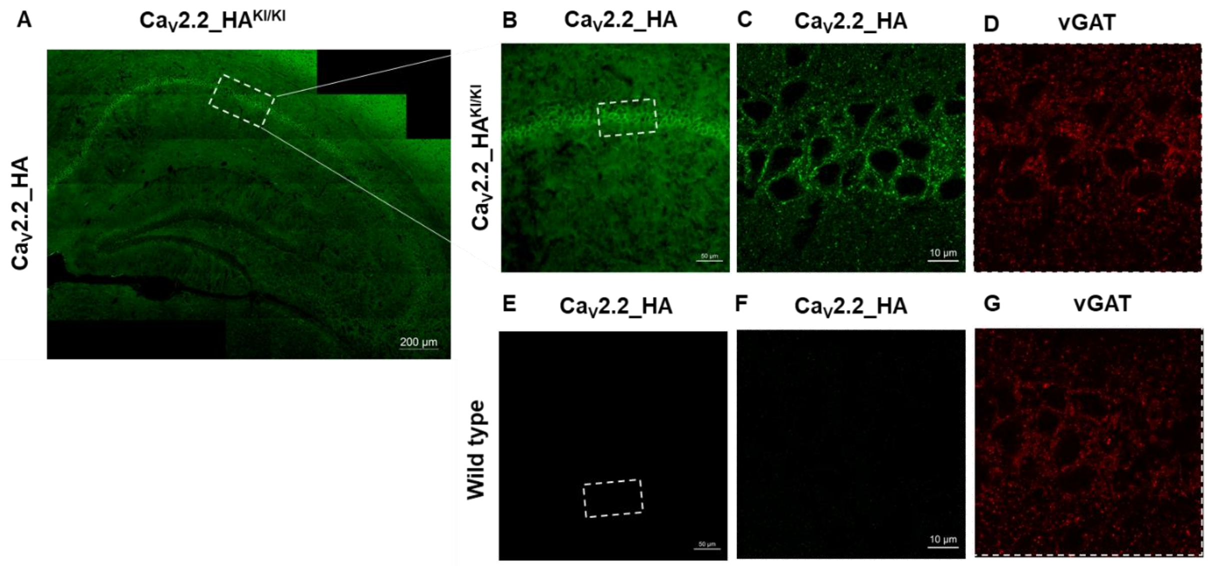


**(A)** Ca$_V$2.2_HA immunolabelling in the hippocampus of adult Ca$_V$2.2_HA$^{KI/KI}$ mice visualised using anti-HA antibodies (green). x20 super resolution tile scan. Scale bar = 200 µm. **(B)** Ca$_V$2.2_HA (green) signal in CA1 area of the hippocampus, x20 confocal imaging, scale bar = 50 µm. **(C)** Ca$_V$2.2_HA (green) x 63 of CA1 somata of Ca$_V$2.2_HA$^{KI/KI}$ mice in super resolution, maximum intensity projection of z-stack with 0.197 µm optical sections, scale bar = 10 µm. **(D)** vGAT signal shown in red in CA1 in super resolution, maximum intensity projection of z-stack with 0.197 µm optical sections, scale bar = 10 µm. **(E-G)** Wild-type control images of Ca$_V$2.2_HA and vGAT at x20 confocal, scale bar = 50 µm (E) and x63 super resolution, maximum intensity projection of z-stack, scale bar = 10 µm (F and G).

**Figure S2 (relates to Fig 4 E). Wild-type control shows lack of $Ca_V2.2$_HA staining in hippocampal neurons.**

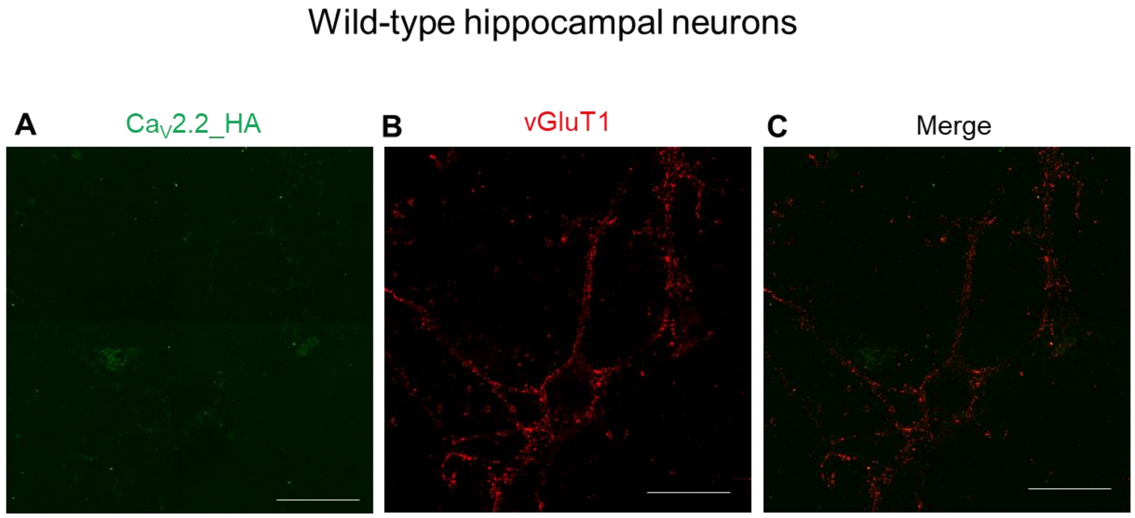


Control Airyscan images for Fig 4 E. In hippocampal neurons from wild-type mice, no staining for $Ca_V2.2$_HA channels was visible **(A)** whereas the vGluT1 **(B,** red) signal was similar to neurons from $Ca_V2.2$_HA$^{KI/KI}$ mice. Panel C shows the merge of the HA and vGluT1 signal. Images were taken with the same settings as the images shown in Fig 4 E. Scale bar = 10 µm.

**Figure S3 (relates to Fig 5 A) Negative no-primary antibody control for $\alpha_2\delta$-1_HA transfected neurons.**

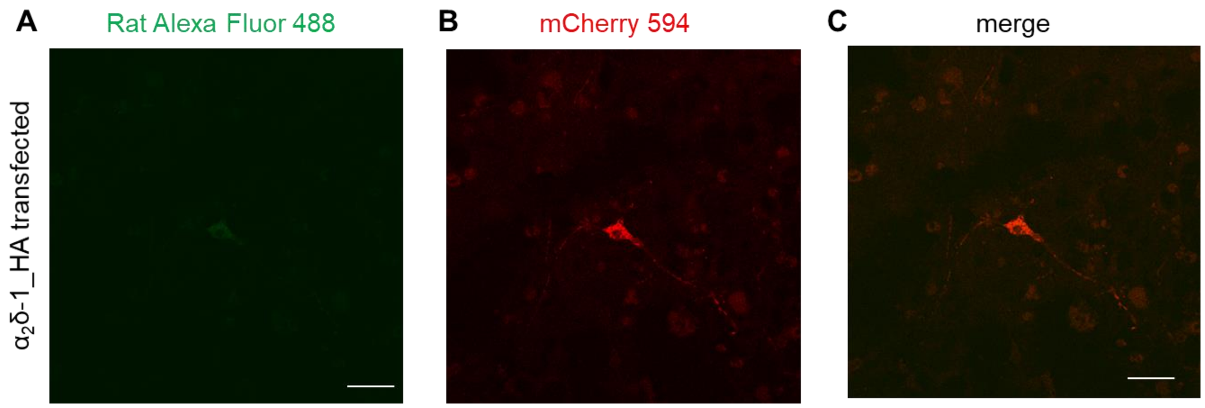


**(A)** Control confocal images for Figure 5 A without anti-HA pABs to label transfected $\alpha_2\delta$-1_HA. **(B)** mCherry positive signal as marker of transfection. **(C)** Merged mCherry and no primary Ab control for $\alpha_2\delta$-1_HA. Images were taken with the same settings as the images shown in Fig 5 A. Scale bar = 5 µm.
